# Rapid Reconstruction of Time-varying Gene Regulatory Networks

**DOI:** 10.1101/272484

**Authors:** Saptarshi Pyne, Alok Ranjan Kumar, Ashish Anand

## Abstract

—Rapid advancements in high-throughput technologies has resulted in genome-scale time series datasets. Uncovering the temporal sequence of gene regulatory events, in the form of time-varying gene regulatory networks (GRNs), demands computationally fast, accurate and scalable algorithms. The existing algorithms can be divided into two categories: ones that are time-intensive and hence unscalable; others that impose structural constraints to become scalable. In this paper, a novel algorithm, namely ‘an algorithm for reconstructing Time-varying Gene regulatory networks with Shortlisted candidate regulators’ (*TGS*), is proposed. *TGS* is time-efficient and does not impose any structural constraints. Moreover, it provides such flexibility and time-efficiency, without losing its accuracy. *TGS* consistently outperforms the state-of-the-art algorithms in true positive detection, on three benchmark synthetic datasets. However, *TGS* does not perform as well in false positive rejection. To mitigate this issue, *TGS+* is proposed. *TGS+* demonstrates competitive false positive rejection power, while maintaining the superior speed and true positive detection power of *TGS*. Nevertheless, main memory requirements of both *TGS* variants grow exponentially with the number of genes, which they tackle by restricting the maximum number of regulators for each gene. Relaxing this restriction remains a challenge as the actual number of regulators is not known a priori.

**Reproducibility:** The datasets and results can be found at: https://github.com/aaiitg-grp/TGS. This manuscript is currently under review. As soon as it is accepted, the source code will be made available at the same link. There are mentions of a ‘supplementary document’ throughout the text. The supplementary document will also be made available after acceptance of the manuscript. If you wish to be notified when the supplementary document and source code are available, kindly send an email to with subject line ‘TGS Source Code: Request for Notification’. The email body can be kept blank.

## 1 Introduction

Cell, the building block of life, is a dynamic system. It continuously senses and responds to the changes in its environmental conditions. The cell performs this task with the help of several biomolecules that interact with each other. Modelling how these interactions vary with time is critical for understanding how the cell develops, evolves and maintains itself. One of the important types of interactions is the ones between the transcription factors (TFs) and the genes. Each TF is a special type of protein which physically binds to an appropriate site in the vicinity of its target gene to regulate the gene’s expression. The gene, that has produced the TF in the first place, is said to be a regulator of the target gene. This regulator-regulatee relationships among the genes is represented by a directed network, known as the gene regulatory network (GRN). In a GRN, each node represents a gene and a directed edge from one gene to another implies that the former gene regulates the latter. The aim of this paper is to develop an algorithm for modelling (reconstructing) how the GRN structure (edge relationships) varies with time.

There exists an array of algorithms ([1], [2], [3], [4], [5], [6], [7], [8], [9], [10]) that reconstruct time-varying GRNs, from time series gene expression datasets. Among them, the *ARTIVA* algorithm [4] demonstrates its ability to accomplish this task with high accuracy. It employs a flexible framework, where time interval specific GRNs are learnt independently of each other, without imposing any structural constraints. However, *ARTIVA* is very time intensive; therefore, not suitable for the contemporary high-throughput datasets.

In this paper, a novel algorithm, namely ‘an algorithm for reconstructing Time-varying Gene regulatory networks with Shortlisted candidate regulators’ (*TGS*), is proposed to provide an equally flexible framework in a significantly more time-efficient manner.

It needs to be noted that there exist more time-efficient algorithms than *ARTIVA*. Most notably, the algorithms, proposed by Dondelinger et al. [5] and Chan et al. [6]. However, they depend upon the smoothly time-varying assumption, which encourages temporally adjacent GRNs to share more common edges than temporally distal GRNs.

The smoothly time-varying assumption does not necessarily hold for all scenarios. For example, temporal variations are highly dynamic, during Yeast’s stress response [4] or clinical interventions [5]. Another exception may arise when sampling interval of the given dataset is considerably large. In that case, two consecutive time points may belong to two different cellular conditions. Thus, it is possible for a GRN to share less common edges with a temporally adjacent GRN, belonging to a separate condition, than with a distal GRN, belonging to a similar condition (see Section 4.1 of the supplementary document).

*TGS*’s framework is compatible with any dataset, regardless of whether the smoothly time-varying assumption holds for it or not. Moreover, *TGS* offers such flexibility and time-efficiency, without losing its accuracy. Three of the datasets, provided by ‘DREAM3 in silico network inference challenge’, are used to evaluate *TGS*’s prediction power, against those of the aforementioned algorithms. The results show that *TGS* is consistently competitive with these state-of-the-art algorithms, in terms of both sensitivity and precision. It is also found that, *TGS* is able to reconstruct biologically meaningful GRNs from a D. melanogaster microarray dataset.

To summarize, the main contribution of this paper is two-fold:

- **Flexibility:** It provides a framework, where time-varying GRN structures are learnt, independently of each other, without imposing any structural constraints. This framework is compatible with any time series gene-expression dataset, regardless of whether the underlying gene regulation process complies with the smoothly time-varying assumption or not.
- **Time-efficiency:** The framework is offered in a significantly more time-efficient manner than those of the state-of-the-art alternatives. At the same time, competitive sensitivity and precision are provided.

This paper is organized as follows: Section 2 formally defines the problem statement at hand. It also provides a critical and chronological survey of the related algorithms, to identify the fundamental issues, that need to be solved. Subsequently in Section 3, a novel algorithm is proposed to resolve a selected subset of these issues. The learning power and speed, of the proposed algorithm, are comparatively studied against those of the prior state-of-the-art algorithms, in Section 4, on a set of benchmark synthetic datasets. In the same section, biological significance of its prediction is evaluated, given a real microarray dataset. Nevertheless, a set of limitations remain in its unconstrained application. These limitations are discussed in Section 5, which opens the door for future investigations.

## 2 Related Works

### 2.1 Problem Formulation

Suppose that the given dataset 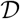 is comprised of *S* time series 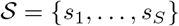 of gene expression data. Each time series contains expression levels of *V* genes 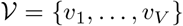 at *T* consecutive time points 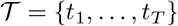. It is assumed that there is no missing value in any time series. In other words, each time series is a complete time series of *T* time points. 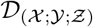 is used to denote the observed values of genes 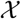 at time points 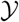 in time series 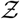. 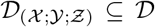 since 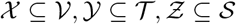.

Given 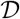, the objective is to reconstruct a temporally ordered GRN sequence 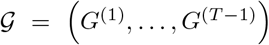 (Figure 1). Here, each 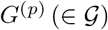 is a time interval specific GRN. Thus, *G*^(*p*)^ represents the gene regulatory events occurred during the time interval between time points *t_p_* and *t*_(*p*+1)_. It is a directed unweighted network on the (2 × *V*) nodes 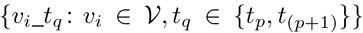. Each node *v_i_*_*t_q_* represents a distinct random variable. Expression of gene *v_i_* at time point *t_q_* is modelled as random variable *v_i_*_*t_q_*. Therefore, the observed expression values of *v_i_* at time point *t_q_* in *S* separate time series are considered as *S* observed values of *v_i_*_*t_q_*. The underlying gene regulation process is assumed to be first order Markovian [1]. Hence, there exists a directed edge (*v_i_*_*t_p_, v_j_*_*t*_(*p*+1)_) if and only if *v_i_*’s expression at time point *t_p_* has a regulatory effect on *v_j_*’s expression at time point *t*_(*p*+1)_.

**Fig. 1.**
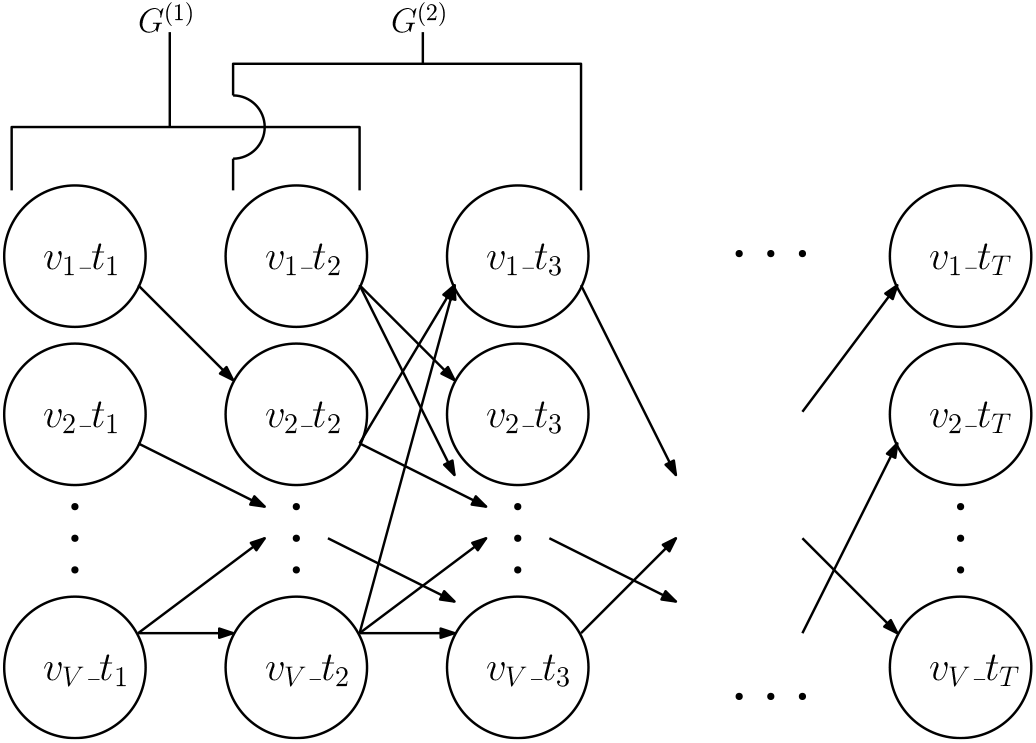
Output time-varying GRNs 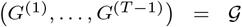 is a sequence of directed unweighted networks. Here, 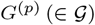 represents the gene regulatory events occurred during the time interval between time points *t_p_* and *t*_(*p*+1)_. It consists of (2 × *V*) nodes 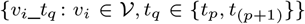. There exists a directed unweighted edge (*v_i_*_*t_p_, v_j_*_*t*_(*p*+1)_) if and only if *v_i_* regulates *v_j_* during time interval (*t_p_, t*_(*p*+1)_).

### 2.2 Existing Solutions

There exists an array of algorithms (e.g., *Bene* [11], *GENIE3* [12], *NARROMI* [13], *LBN* [14]) that do not completely solve the problem but can reconstruct a single ‘summary’ GRN over the nodes in 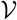. These algorithms consider 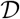 as a cross-sectional dataset. Then they include edge (*v_i_, v_j_*) in the summary GRN if and only if the expression level of *v_j_* is not conditionally independent of that of *v_i_*, given the expression levels of other genes. The summary GRN helps to discover which gene regulates which gene. However, it can not identify the time interval(s) during which such regulation has taken place.

To address this problem, Friedman et al. [1] model 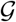 as a Dynamic Probabilistic Network (or Dynamic Bayesian Network, in short, *DBN*). It is assumed that each gene is regulated by the same regulator(s), if any, at every time interval. Thus, output time-varying GRNs possess time-homogeneous structures [Section 1] [15], e.g., edge (*v_i_*_*t*_1_, *v_j_*_*t*_2_) exists in *G*^(1)^ if and only if edge (*v_i_*_*t*_2_, *v_j_*_*t*_3_) exists in *G*^(2)^ and so on [1]. In practice, this approach tends to discover the regulations that are active over a large number of time intervals. However, they may miss the regulations that are active over a small number of time intervals [16].

To tackle the time-homogeneity issue, four algorithms are proposed. They are Non-stationary dynamic Bayesian networks (*NsDbn*) [2], Non-stationary continuous dynamic Bayesian networks (*NsCdbn*) [3], *ARTIVA* [4] and *cpBGe* [15]. These algorithms accommodate the case where the same gene can be regulated by different regulators during different time intervals. The class of resulting models is known as time-inhomogeneous DBN. *NsDbn* assumes that the data is generated by a multiple change point process with change points at 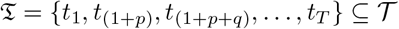 where *p, q* ∈ ℤ_+_. The duration between two consecutive change points is called a time segment. One GRN structure is learnt for each time segment instead of each time interval. GRN structures of two different time segments can be different but GRN structure within a time segment remains unchanged. First, *NsDbn* determines the number of the change points along with their positions. Then for each time segment, the fittest (w.r.t. a scoring function) DBN structure is identified to be the output GRN for that segment. The scoring function requires the input dataset to be discretized. In contrast, *NsCdbn* uses a different scoring function that does not require the dataset to be discretized [3].

*ARTIVA* provides a more flexible model by relaxing the assumption that all genes share the same change points. It assumes that each gene has its own set of change points (i.e. time segments specific to itself); therefore, it can be regulated by different regulators at its different time segments but within a time segment, it must be regulated by the same regulators. Hence, the output is one GRN structure for each time interval. If a particular time segment of a specific gene spans multiple consecutive time intervals, then the gene’s regulator configurations (incoming edges) remain the same in the corresponding time interval specific GRN structures. On the other hand, if two consecutive time intervals belong to two separate time segments for a specific gene, then the gene’s regulator configurations vary in the corresponding time interval specific GRN structures. Thus, *ARTIVA* learns the time interval specific GRN structures independently of each other [4].

Grzegorczyk et al. [15] argue that the assumption of *NsDbn* is over-restrictive (same change points for all the genes) and that of *ARTIVA* is over-flexible (unique change points for every gene) [15]. Being true to their argument, they propose the *cpBGe* algorithm. It groups the genes into clusters based on similar expression patterns and then infers unique change points for every cluster. However, *cpBGe* produces a single time-invariant GRN structure. Therefore, *ARTIVA* remains the most viable alternative for reconstructing time-varying GRNs till that point.

In line with Grzegorczyk et al., Dondelinger et al. [5] argue that the flexibility of *ARTIVA* may lead to overfitting, when *S* ≪ *V*. Hence, they propose an alternative framework, that allows ‘Information sharing’ or ‘coupling’ between time interval specific GRN estimators. The framework is further categorized into two classes: hard coupling and soft coupling, depending upon the strength of coupling (expected similarity between time interval specific GRNs). For the hard coupling, two algorithms, *TVDBN-bino-hard* and *TVDBN-exp-hard*, are introduced; they assume that the expression of each gene follows a binomial and an exponential distribution, respectively. Similarly, for the soft coupling, two more algorithms, *TVDBN-bino-soft* and *TVDBN-exp-soft*, are proposed. Dondelinger et al. also propose an unconstrained (no ‘Information sharing’) variant, called *TVDBN-0*, which is same as *ARTIVA*, except in the internal sampling strategies [5]. Through a comparative study, over a collection of simulated datasets, it is concluded that ‘Information sharing’ improves prediction when the true network varies smoothly with time (each GRN structure shares more common edges with its temporally adjacent GRN structures than with the distal ones, known as the ‘smoothly time-varying assumption’). Following the same assumption, Chan et al. [6] propose a time-efficient algorithm (henceforth, *MAP-TV*), based on a Maximum A Posteriori (MAP) probability estimation approach. Later, Zhang et al. [10] replace the *L*_1_-based penalties in *MAP-TV* with a multi-Laplacian prior. This approach further helps in simultaneously obtaining time-varying GRNs and transcriptional regulatory networks.

Nevertheless, it is always preferable to avoid any structural assumptions, unless the assumptions are known a priori to hold for the dataset [4]. *ARTIVA* offers such an unconstrained framework. It learns each time interval specific GRN structure in a purely data-driven manner, independently of other GRN structures.

However, *ARTIVA* is a computationally expensive algorithm. Its authors suggest that it requires approximately (5 × *V*) minutes to reconstruct GRNs from a dataset of *T* = 20 and *S* = 5 with the default parameter settings on a 2.66 GHz CPU having a 4 GB RAM [4]. Hence, the time frame necessary for it to scale up to the high-throughput datasets may be considered prohibitive (≃ 87 days for *V* = 25, 000). Developing an unconstrained as well as time-efficient framework can be considered a timely contribution. A summary of the algorithms, discussed in this section, is provided in Table 1.

**TABLE 1.**
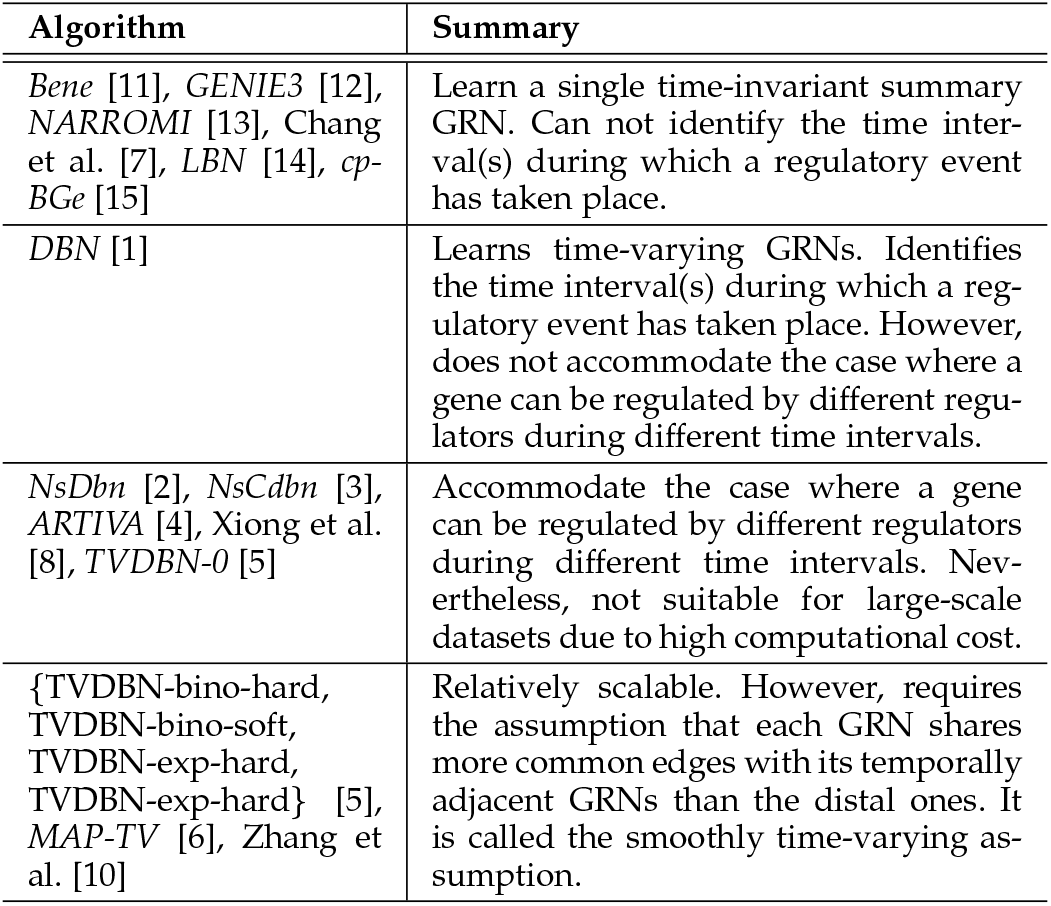
A Summary of the existing GRN Reconstruction Algorithms, discussed in Section 2.2, in the context of Time Series Gene Expression Data.

| Algorithm | Summary |
| --- | --- |
| *Bene* [11], *GENIE3* [12], *NARROMI* [13], Chang et al. [7], *LBN* [14], *cp-BGe* [15] | Learn a single time-invariant summary GRN. Can not identify the time interval(s) during which a regulatory event has taken place. |
| *DBN* [1] | Learns time-varying GRNs. Identifies the time interval(s) during which a regulatory event has taken place. However, does not accommodate the case where a gene can be regulated by different regulators during different time intervals. |
| *NsDbn* [2], *NsCdbn* [3], *ARTIVA* [4], Xiong et al. [8], *TVDBN-0* [5] | Accommodate the case where a gene can be regulated by different regulators during different time intervals. Nevertheless, not suitable for large-scale datasets due to high computational cost. |
| {*TVDBN-bino-hard*, *TVDBN-bino-soft*, *TVDBN-exp-hard*, *TVDBN-exp-soft*} [5], *MAP-TV* [6], Zhang et al. [10] | Relatively scalable. However, requires the assumption that each GRN shares more common edges with its temporally adjacent GRNs than the distal ones. It is called the smoothly time-varying assumption. |

## 3 Methods

In this section, two algorithms are developed. First, a baseline algorithm is developed for reconstructing time-varying GRNs from time series gene expression data. Like ARTIVA [4], the baseline algorithm attempts to reconstruct the time-varying GRNs independently of each other. Therefore, it is compatible with any dataset, regardless of whether the smoothly time-varying assumption holds for it or not. Nevertheless, it is time-intensive and hence not suitable for large-scale datasets. Second, a heuristic based approximation step is added to the baseline algorithm to develop the final algorithm. It maintains the independently time-varying framework without compromising the scalability.

### 3.1 Development of the Baseline Algorithm

A conditional independence based baseline algorithm, referred to as Time-varying Bayesian Networks (*TBN*), is designed. Algorithm 1 describes the steps in *TBN*. It takes a discretized complete time series gene expression dataset 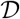 as input. It is assumed that there are multiple time series (*S* > 1) in 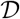. Then *TBN* reconstructs one GRN *G*^(*p*)^ for every time interval (*t_p_, t*_(*p*+1)_), where 1 ≤ *p* ≤ (*T* − 1) (Figure 1). Each *G*^(*p*)^ is modelled as a Bayesian Network (BN) [17]. Absence of a directed edge (*v_i_*_*t_p_, v_j_*_*t*_(*p*+1)_) in *G*^(*t_p_*)^ implies that the expression level of *v_j_* at time point *t*_(*p*+1)_ is conditionally independent of that of *v_i_* at time point *t_p_*, given the expression levels of the genes 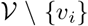 at time point *t_p_*. Biologically, it signifies that the expression level of *v_i_* at time point *t_p_* has no regulatory effect on that of *v_j_* during time interval (*t_p_, t_p_* + 1). On the other hand, presence of that edge signifies that there is a nonzero probability that *v_i_*’s expression level at *t_p_* has affected that of *v_j_* during the (*t_p_, t_p_* + 1) time interval.

*TBN* employs a BN structure learning algorithm [18] to learn every *G^(p)^* from 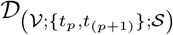. Therefore, the problem of learning (*T* − 1) time-varying GRNs in 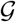 gets decomposed into (*T* − 1) independent BN structure learning problems. For learning an exact BN structure, *Bene* is the state-of-the-art algorithm w.r.t. time complexity and scalability, to the best of authors’ knowledge. Hence, *TBN* with *Bene* is chosen as the baseline for developing a novel algorithm.

In *TBN* (Algorithm 1 Line 17), BIC scoring function [17] is used with *Bene* to compute scores of the candidate regulator sets. There exist some other scoring functions that can be used with *Bene*, e.g. BDe. Among all available scoring functions, BIC and BDe are compared w.r.t. their effects on learning power of *Bene* by Silander et al. [11]. It is observed that BIC outperforms BDe when number of observations being considered (here, (*S* + 1)) is below 20. Moreover, the performance of BDe is very sensitive to the chosen value of its hyper-parameter [19]. BIC, on the other hand, does not depend on any hyper-parameter. For these reasons, BIC is considered to be most suitable for the current study.

### 3.2 Development of a Novel Algorithm: ‘an algorithm for reconstructing Time-varying Gene regulatory networks with Shortlisted candidate regulators’ (TGS)

From Algorithm 1, it is found that TBN’s time complexity is *T*_TBN_ (*V*) = (*T* − 1) × *V* × *o*(*V*^2^2^(*V*−2)^) = *o*((*T* − 1) *V*^3^2^(*V*−2)^). It grows exponentially with the number of candidate regulators for each gene, which is *V* in this case. Therefore, this approach can be made more computationally efficient if a way can be discovered that: (a) generates a significantly shorter list of candidate regulators for each gene, and (b) the amount of time it spends for shortlisting candidate regulators is overshadowed by the time gain it brings.

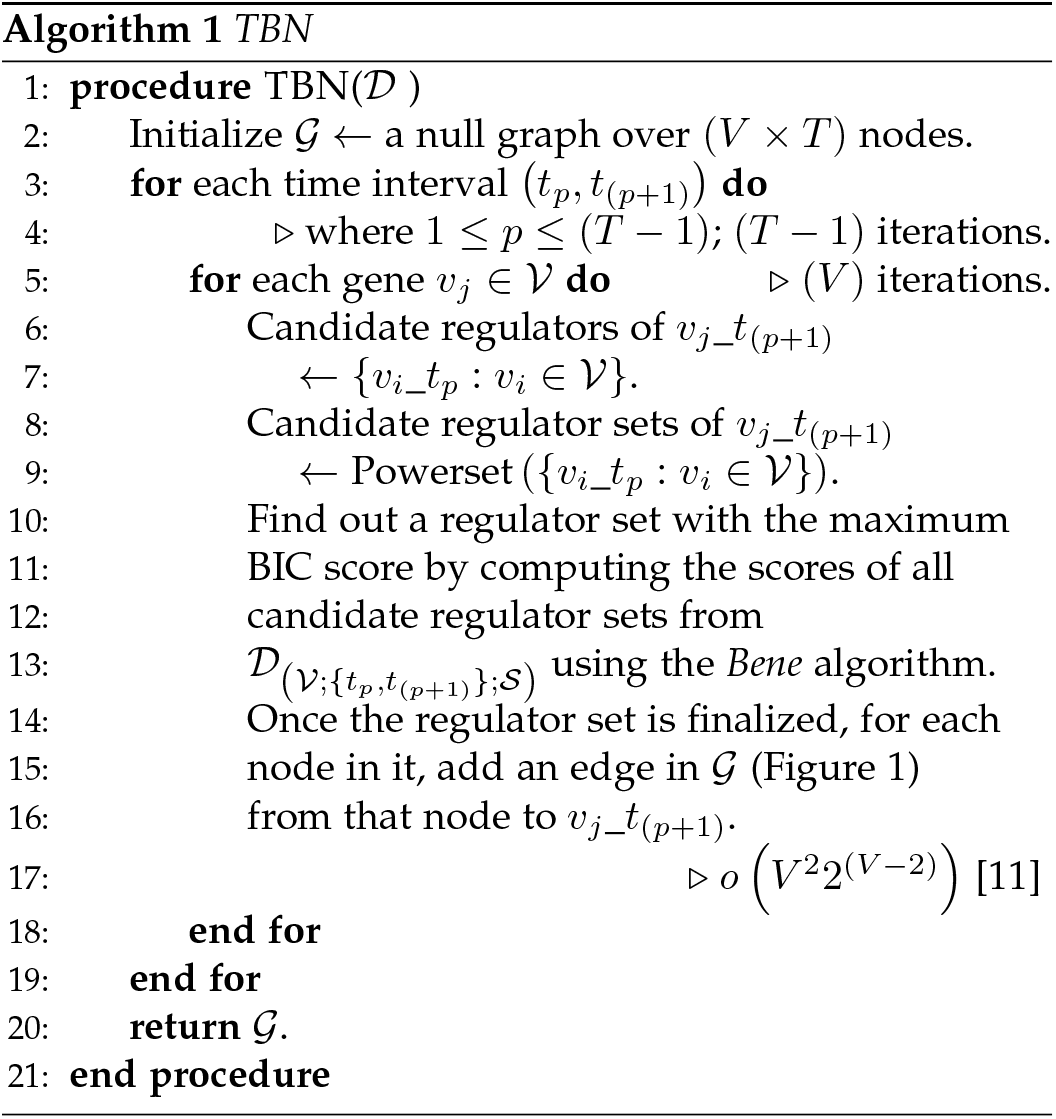

Statistical pairwise association measures fulfil the first criterion. Given sufficient observations on a pair of random variables, they can identify whether there is a statistically significant probability (w.r.t. a predefined significance threshold) that these variables are not associated with each other. Thus the candidate regulators, whose expressions are not statistically associated with that of the regulatee gene, could be identified. Then these regulators can be removed from the candidate regulator set.

A study by Liu et al. [20] compares 14 such association measures and concludes that Mutual Information (MI) demonstrates superior stability over other measures. MI’s potential regulator-regulatee association predictions consistently outperform [20] those of most others across different sizes (different values of *V*) of benchmark gene expression datasets w.r.t. mean AUC (Area Under Receiver Operating Characteristic Curve). Algorithms *NARROMI* and *LBN* utilize MI for short-listing candidate regulators. For each regulatee gene, they calculate its MI with every candidate regulator; then eliminate the candidates with MI lower than a user defined threshold.

However, the Achilles’ heel of this strategy is that the prediction is heavily dependent on the user defined threshold value [14]. *LBN* determines the threshold for synthetic datasets by performing the predictions multiple times with different threshold values and choosing the one that gives the best prediction. This threshold selection strategy requires the true regulatory relationships to be known a priori so that quality of a prediction can be measured. A more practical strategy is appointed by Context Likelihood of Relatedness (*CLR*) algorithm [21] (Algorithm 2). It constructs a weighted MI network^1^ over all genes from a gene expression dataset without requiring a user defined threshold. *CLR* is found to outperform other major MI network inference algorithms [21]. Furthermore, it requires only 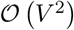 time for a dataset with *V* genes.

For the aforementioned reasons, *CLR* is chosen to be a pre-selection step for candidate regulators before more comprehensive selection could be performed by *TBN*. It gives birth to a novel algorithm, which is named *TGS* (short form for ‘the algorithm for reconstructing Time-varying Gene regulatory networks with Shortlisted candidate regulators’) (Algorithm 3). A flowchart is presented in Section 4.2 of the supplementary document. Additionally, a graphical flowchart, with a small example, is depicted in Figure 2.

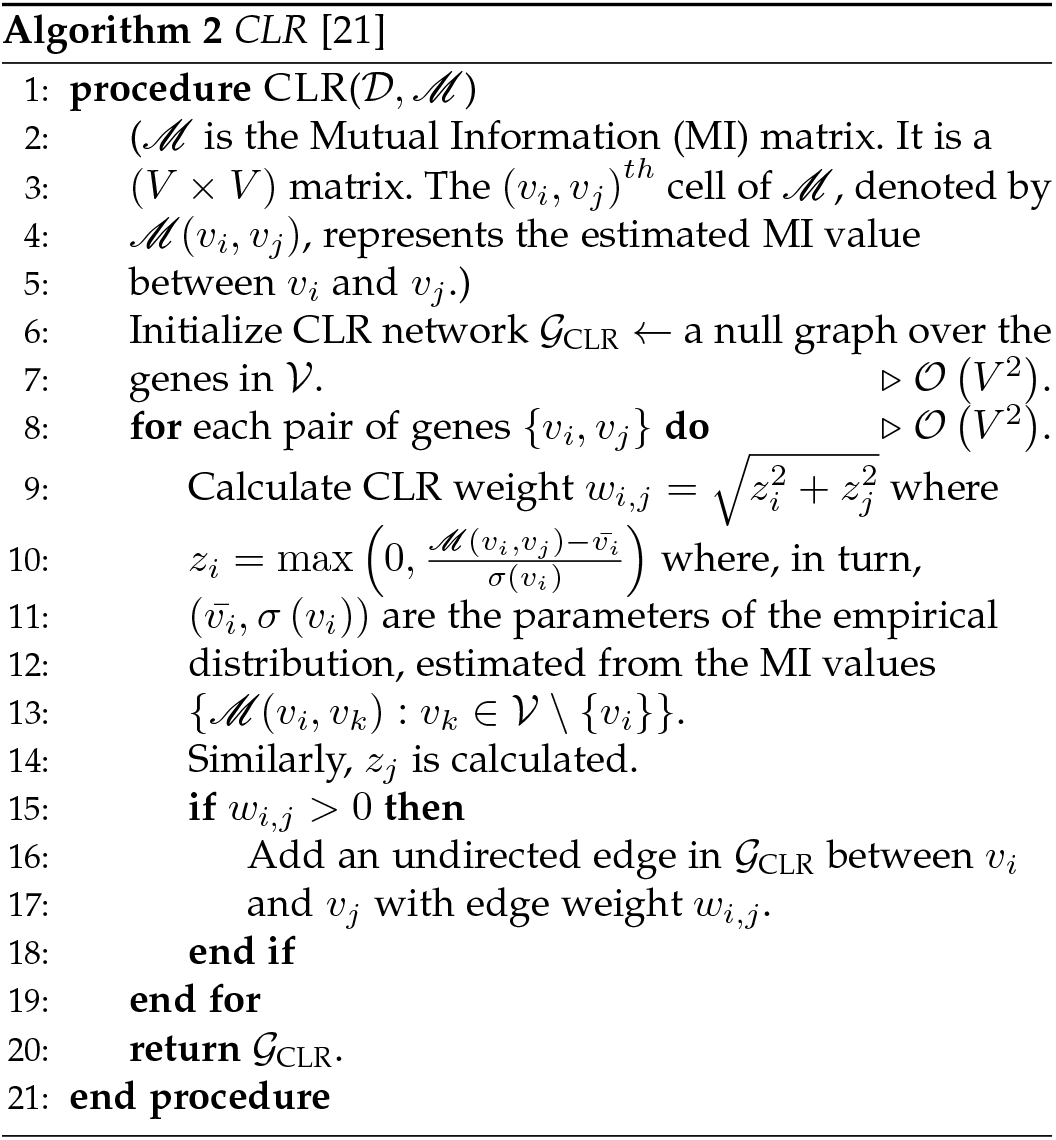

**Fig. 2.**
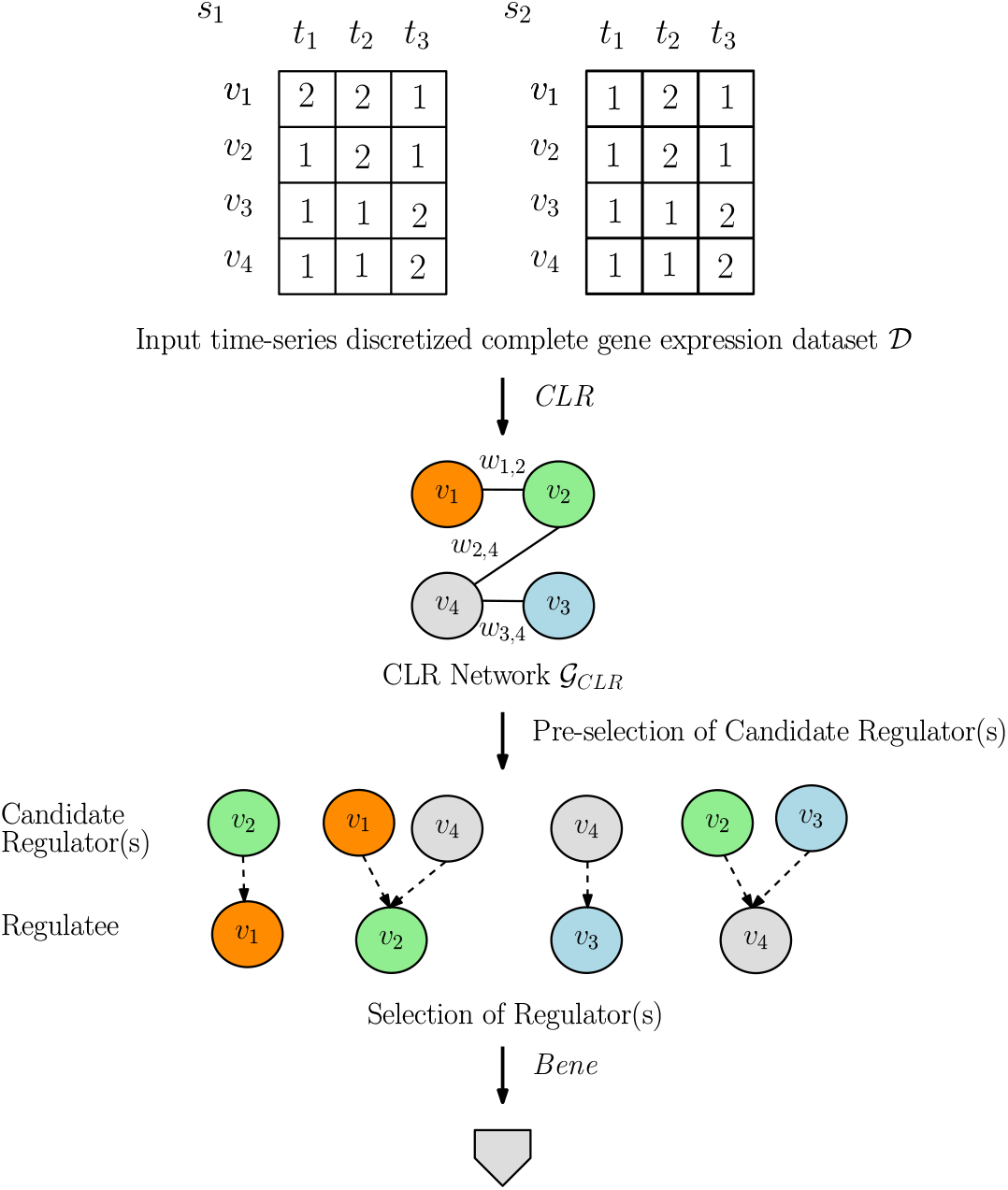
Graphical Flowchart (Part 1) of the *TGS* Algorithm (Algorithm 3). The flowchart is continued in Figure 3. For illustration, a dataset 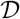 is considered with four genes 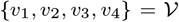 and two time series 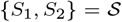. Each time series has three time points 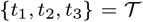. 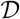 is discretized into two discrete levels, represented by {1, 2}.

TGS (Algorithm 3) has the time complexity 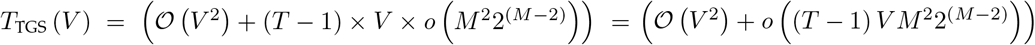. Here, *M* is the maximum number of neighbours a gene has in the *CLR* network. Since, in theory, *M* ≤ *V*, time complexity of *TGS* is upper bounded by that of *TBN*. But empirically, it is found [22] that each gene is regulated by a small number of regulators with the exception in case of E.coli. For this reason, major BN based algorithms (e.g., the *DBN* implementation in BayesNet Toolbox for MATLAB [23]) have variants that allow the user to specify the maximum number of regulators a gene can have in a given dataset, known as the max fan-in value (*M_f_*). For each gene, it reduces the number of candidate regulator sets from 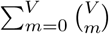 to 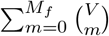, where *M_f_* ≪ *V*. It is further reduced to 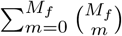 in the variant of *TGS* with max fan-in restriction (Algorithm 4). Therefore, for a high-throughput human-genome scale time series gene expression dataset where (*T* − 1) = *o*(*V*) and *M_f_* = *o*(lg *V*), the time complexity of *TGS* asymptotically tends towards polynomial while that of *TBN* remains exponential.

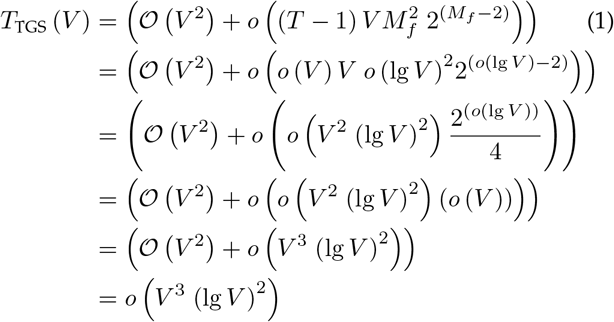

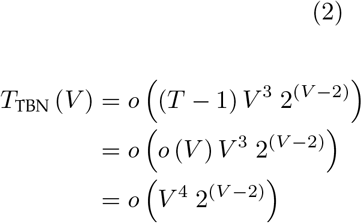

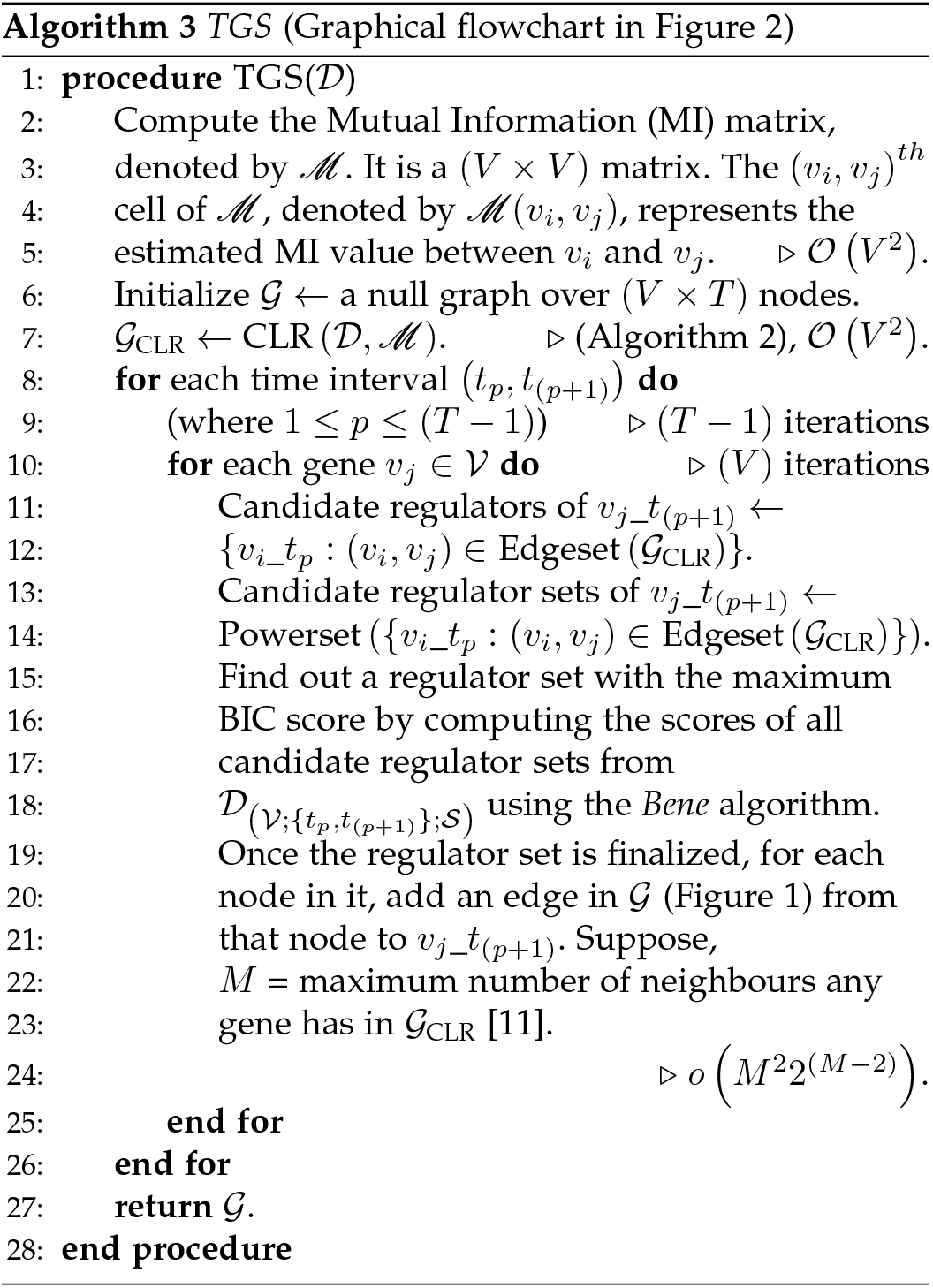

**TGS+:** A variant of Algorithm 4, called *TGS*+, is also proposed to minimize the negative effect of the *CLR* step on false positive rejection. While *CLR* is very beneficial for predicting a useful number of true positive edges, it can do so at the cost of predicting a considerable number of false positive edges [21]. Having such a considerable number of false positives could result in the waste of scarce and often expensive resources in the validation process.

Cases where CLR’s judgment fails are when two genes, *v_i_* and *v_j_*, have a high mutual information, 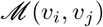, even though they do not have a direct regulatory relationship. One such case is when *v_i_* and *v_j_* have an indirect regulatory relationship. For example, *v_i_* regulates a third gene *v_k_* and *v_k_*, in turn, regulates *v_j_*.

The Data Processing Inequality (DPI) [24] helps to identify the false positive edge in the aforementioned case. The inequality states that the mutual information of the false positive pair must not be higher than those of the true positive pairs: 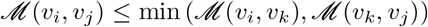. An algorithm, namely *ARACNE* [24], applies the converse of the DPI to produce significantly low number of false positives [24]. *ARACNE* takes the raw mutual information matrix as input. Then, for each 3-combination of genes, it identifies the pair with the lowest mutual information and replace their mutual information with zero in the mutual information matrix (Algorithm 5). Thus, *ARACNE* refines the raw mutual information matrix.

In TGS+ (Algorithm 6), the refined mutual information matrix, instead of the raw one, is provided as input to the *CLR* step. The expectation is that the reduction in false positives in the mutual information matrix will help *CLR* to make less false positive predictions. If that becomes the case, then *TGS+* is expected to predict less false positives, compared to *TGS*.

**Fig. 3.**
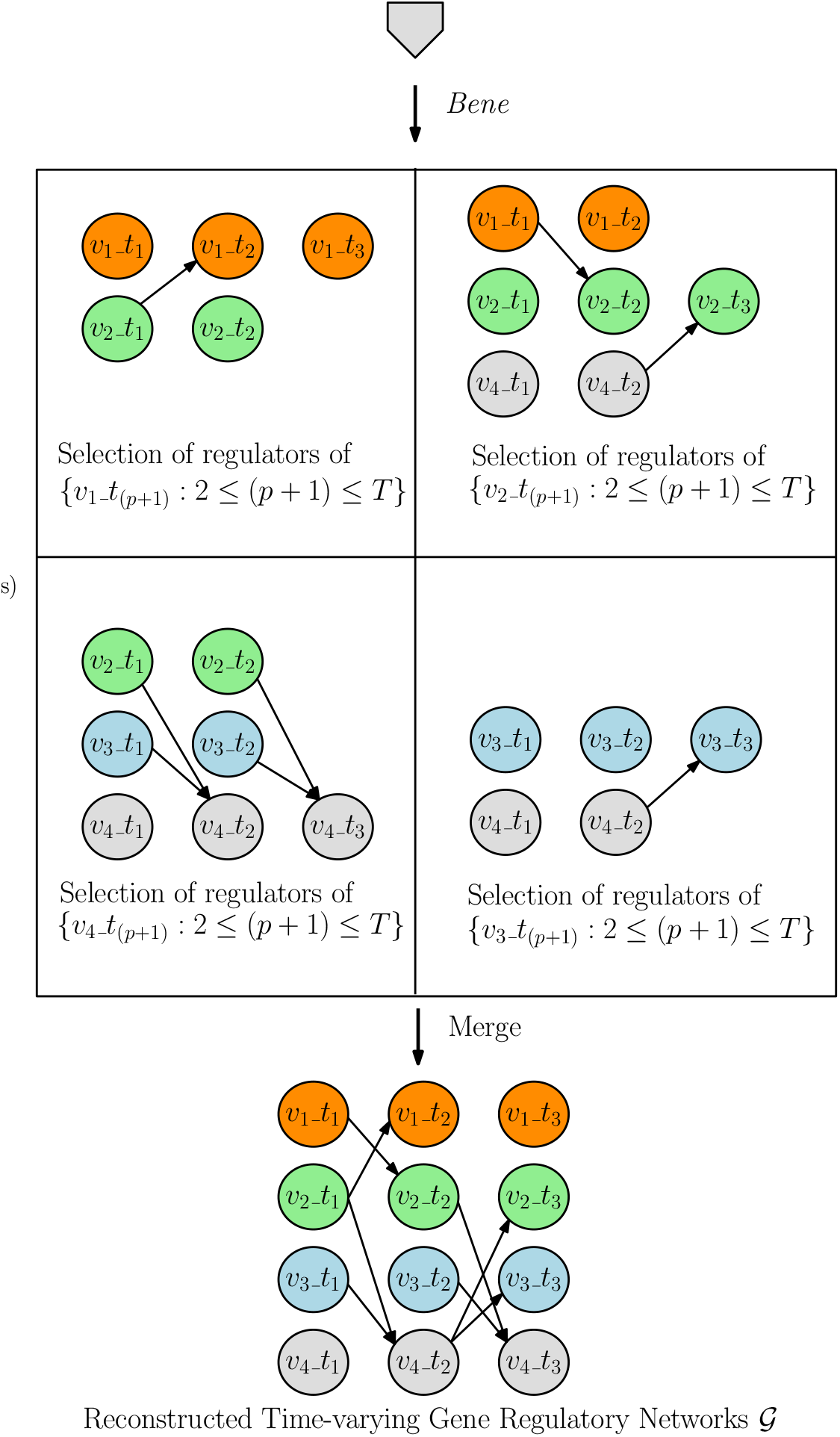
Graphical Flowchart (Part 2) of the *TGS* Algorithm (Algorithm 3). The flowchart is continued from Figure 2. For discussion of the *Bene* step, let us consider the ‘Selection of regulators of {*v*_1__*t*_(*p*+1)_: 2 ≤ (*p* + 1) ≤ *T*}’. Since, *v*_2_ is the sole neighbour of *v*_1_ in *G*_CLR_, *v*_2_ is the only candidate regulator of *v*_1_ (Figure 2). Therefore, the candidate regulator sets of *v*_1__*t*_2_ are 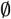 and {*v*_2__*t*_1_}. Among these two sets, *Bene* chooses {*v*_2__*t*_1_} based on observations 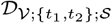. Similarly, the candidate regulator sets of *v*_1__*t*_3_ are 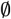 and {*v*_2__*t*_2_}. Among these two sets, *Bene* chooses 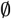 based on observations 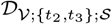.

Nevertheless, *ARACNE*’s power of false positive rejection comes at a cost. When feed-forward network motifs (e.g., *v_i_* regulates *v_j_* directly as well as indirectly through *v_k_*) are present in the true network, *ARACNE* may reject the true feed-forward edges. Another case, where *ARACNE* may result in rejection of the true positive edges, is discussed in Section 4.3 of the supplementary document. However, it is observed that the improvement in false positives, due to *ARACNE*, outweighs the decline in true positives [24].

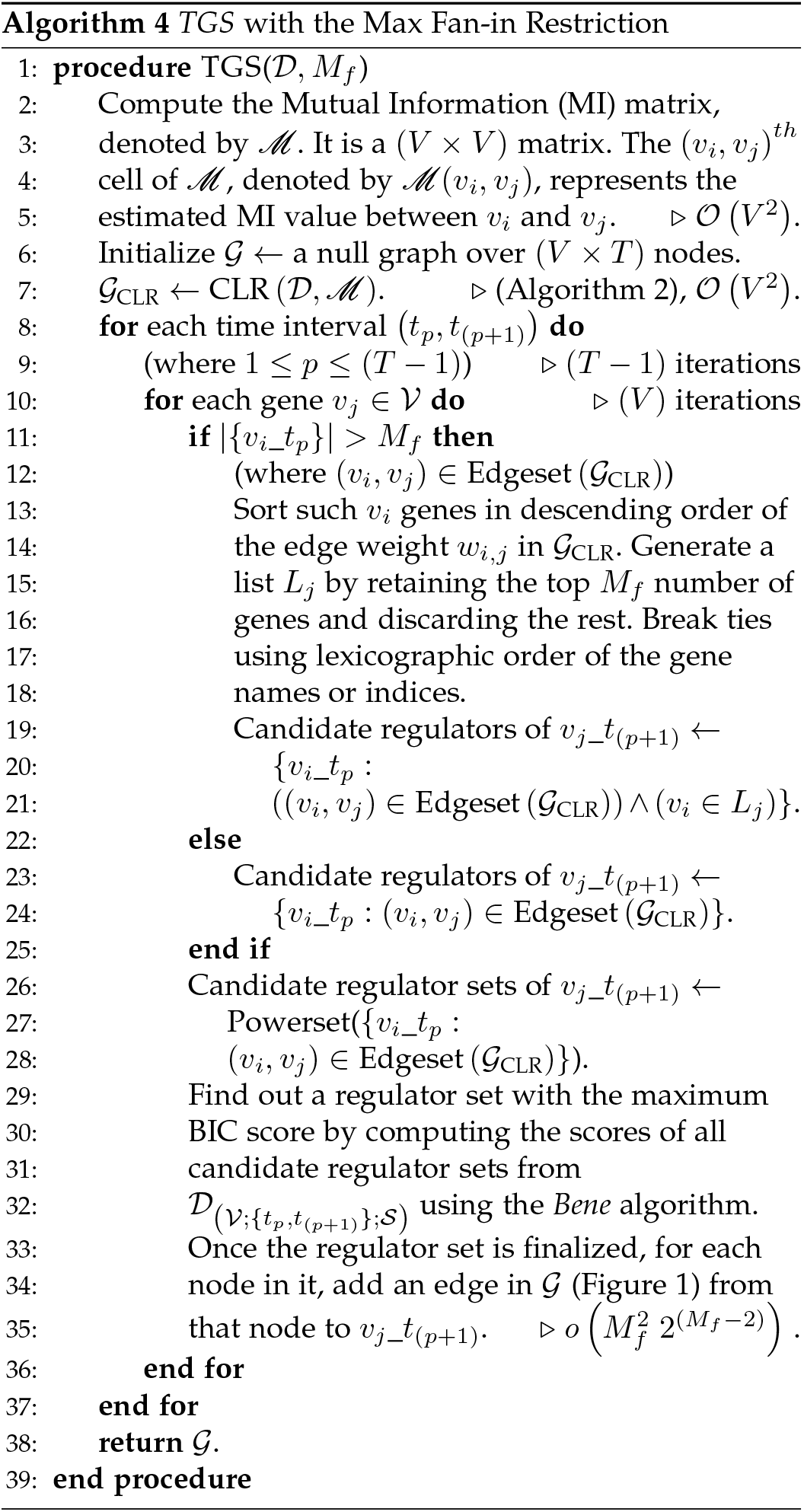

Therefore, *TGS+* is expected to provide a significantly less number of false positives at the loss of slightly less number of true positives, compared to *TGS*. Moreover, since the refined mutual information matrix tend to be sparser than the raw one, *CLR* is expected to shortlist a smaller number of candidate regulators, for each regulatee gene. As a consequence, *TGS+* is expected to have a shorter runtime than even *TGS*. The time complexity of *TGS+* is presented at Section 4.4 of the supplementary document. The differences between *TGS* and *TGS+* are presented as black box diagrams in Figure 4.

**Fig. 4.**
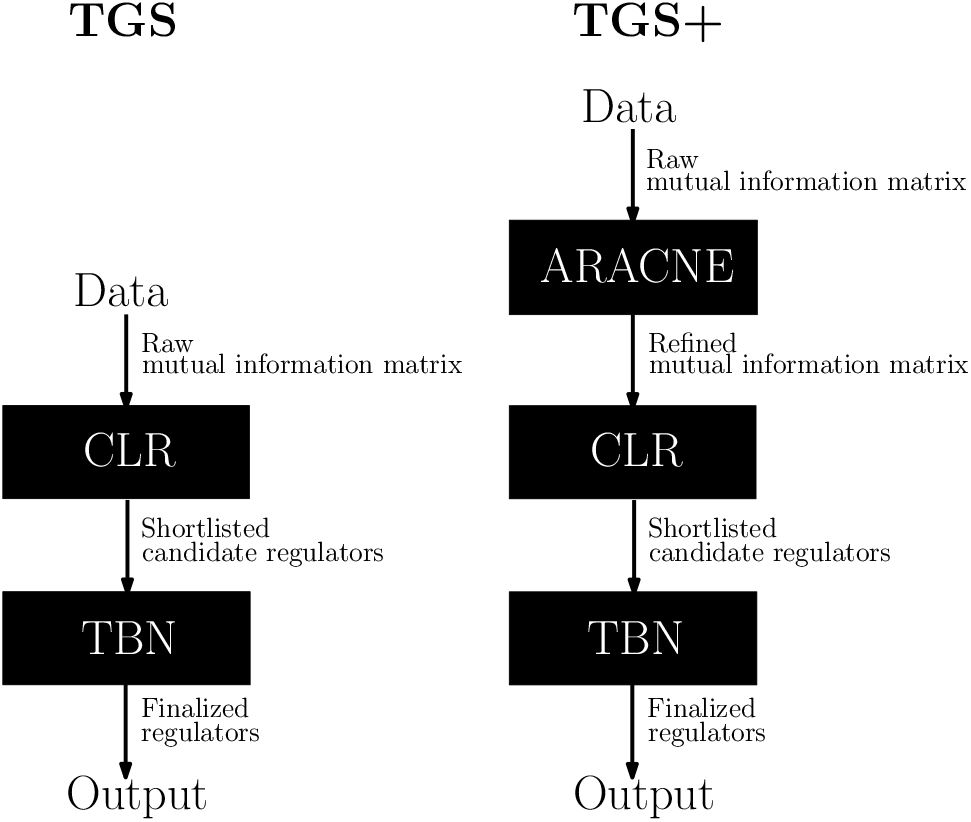
The Black Box Diagrams of the *TGS* and *TGS+* Algorithms. *TGS+* has an additional step, namely *ARACNE*, compared to *TGS*.

## 4 Results

The results of the *TGS* and *TGS+* algorithms, on a set of realistically simulated (synthetic) benchmark datasets and a real microarray dataset, are presented in this section. The learning power and speed of *TGS* and *TGS+* are evaluated with the synthetic datasets, against those of *ARTIVA*, *TVDBN-0, TVDBN-bino-hard* and *TVDBN-bino-soft*. The reason why *TVDBN-exp-hard* and *TVDBN-exp-soft* can not be included in this comparative study, is explained in a footnote of Section 4.3. Moreover, the binomial variants are observed to provide more consistent predictions than the exponential variants [5]. Therefore, the exclusion of the exponential variants, may not be disadvantageous for this study. The *MAP-TV* algorithm can not be included in this study, either, as its source code is not available. An indirect comparative study, between MAP-TV and the *TGS* variants, are presented in Section 4.5 of the supplementary document. Finally, *TGS* and *TGS+* are applied on the real dataset, and biological significance of their predictions are evaluated against the existing biological knowledge.

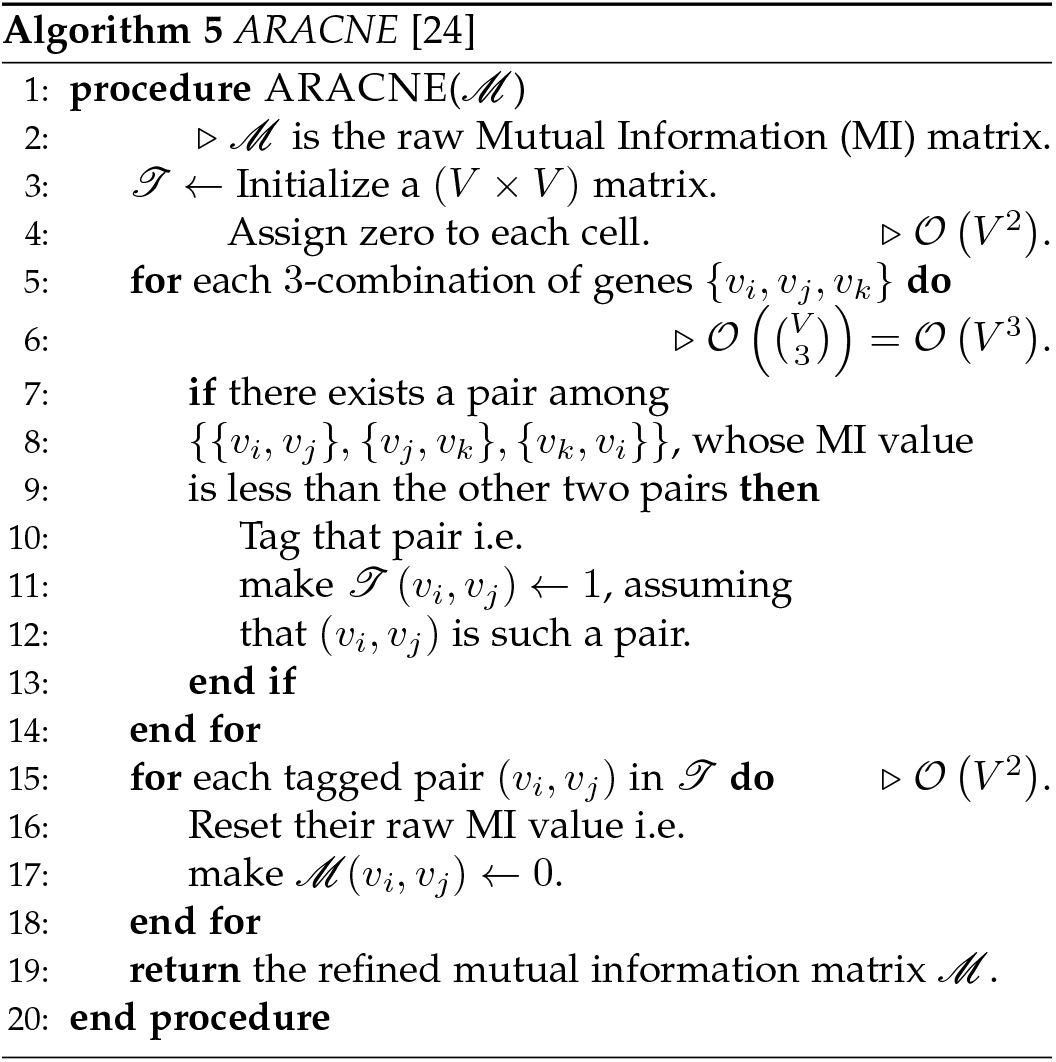

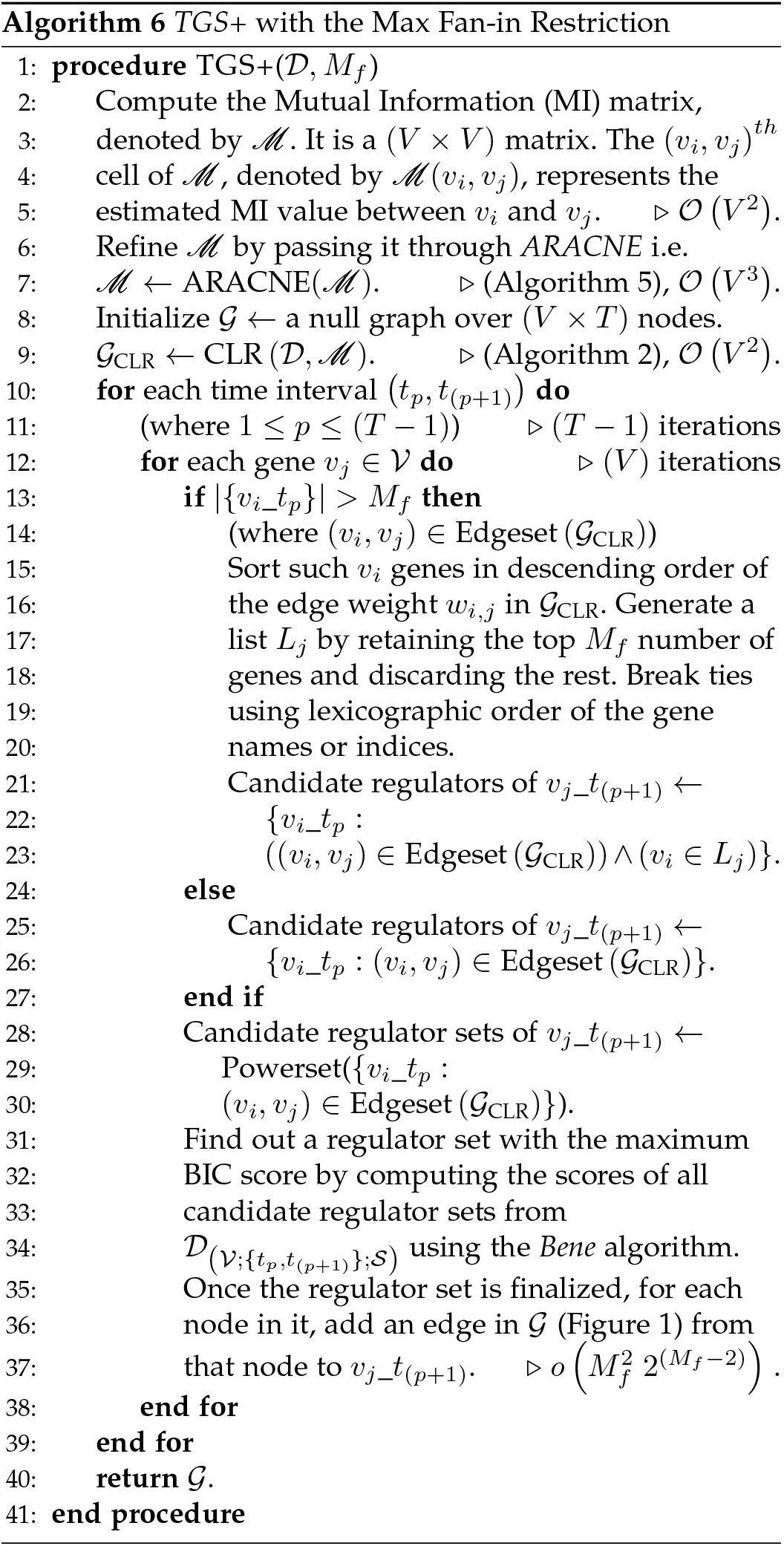

### 4.1 Datasets

#### 4.1.1 Synthetic DREAM3 In Silico Network Inference Challenge Datasets

A real gene expression dataset with known true underlying GRNs is the coveted choice of dataset for evaluating GRN modelling algorithms. To that end, Marbach et al. [25] design three sets of realistic GRN structures for different model organisms with 10, 50 and 100 genes. Then for each of these in silico GRNs, they choose an appropriate dynamical model and generate a dataset through simulation [26]. These datasets are made publicly available as benchmarks for assessing and comparing the modelling algorithms through DREAM3 In Silico Network Challenge [27], [28]. In each dataset, gene expressions are normalized so that the maximum gene expression value in a data file is one. Among these datasets, the ‘Yeast1’ time series datasets are chosen for the purpose of this paper. It comprises of two sets of datasets: one noiseless and the other noisy (resulted from adding Gaussian noise to the noiseless datasets). Each set contains three datasets as summarized in Table 2. It can be noted that the true networks are single network GRNs.

**TABLE 2.** A Summary of the chosen DREAM3 Datasets. Here, *V* = number of genes, *T* = number of time points and *S* = number of time series in a given dataset.

| Datasets (noiseless, noisy) | V | T | S | No. of True Edges |
| --- | --- | --- | --- | --- |
| (Ds10, Ds10n) | 10 | 21 | 4 | 10 |
| (Ds50, Ds50n) | 50 | 21 | 23 | 77 |
| (Ds100, Ds100n) | 100 | 21 | 46 | 166 |

The noisy datasets are chosen for the comparative study because: (a) the presence of noise makes them more realistic than the noiseless datasets, and (b) the noisy datasets are used to evaluate algorithms in the DREAM challenge, while the noiseless datasets are released after the challenge; therefore, the reader can compare the performances of the algorithms, in this study, with those of the algorithms, employed during the challenge.

#### 4.1.2 Real Drosophila melanogaster (Dm) Life Cycle Dataset (DmLc)

The DmLc dataset is a time series real gene expression dataset of the fruit fly’s developmental cycle (from embryonic stage up to the first thirty days of adulthood). This dataset is experimentally produced by Arbeitman et al. [29]. In Song et al. [30], the dataset is utilized for evaluating performance of the *KELLER* algorithm. Since, the original dataset is real-valued and *KELLER* requires a discretized dataset, the original dataset is discretized using the *2L.Tesla* algorithm (Algorithm 2, Section 4.6, supplementary document). The discretized DmLc dataset is obtained from the *KELLER* website [31]. It contains a single time series (*S* = 1) observations of 4028 (*V* = 4028) genes which represent around one-third of identified Dm genes [29]. There are a total of 66 time points (*T* = 66) in the time series, spread across four life cycle stages; they are the Embryonic (E; time points 1-30), Larval (L; time points 31-40), Pupal (P; time points 41-58) and Adult (A; time points 59-66) stages. The sampling intervals vary between different stages and even within a stage. For example, the sampling intervals are either 0.5 hour or 1 hour in the E stage; whereas, it is in order of days in A stage. This is done because the developmental changes happen at different rates in different stages, like - in the E stage, changes take place faster than those in the A stage. The sampling intervals are described in Arbeitman et al. [29]. Following Song et al. [30], a subdataset (hereafter, DmLc3) is also generated from the DmLc dataset by selecting the data corresponding to only a subset of 588 genes, known to be involved in the developmental process of Dm according to their gene ontology annotations. Then the DmLc3 dataset is further divided into four mutually exclusive and collectively exhaustive sub-datasets with respect to four separate stages. These datasets are appropriately named as DmLc3E, DmLc3L, DmLc3P and DmLcA.

### 4.2 Discretization of Data for {*TBN, TGS, TGS+*}

Two different algorithms are chosen for data discretization: one based on domain-knowledge (the wild type values of the genes); another based on a domain-independent strategy. It is observed that the domain-knowledge based algorithm improves learning over the domain-independent one (Section 4.6, supplementary document). Therefore, for all the experiments reported here, the former algorithm is applied to discretize data.

### 4.3 Implementation

*TBN, TGS* and *TGS+* are implemented in `R` programming language [32] version 3.3.2. For *ARACNE* and *CLR*, their implementations in `R` package minet [33] (version: 3.34.0) are used. `R` package bnstruct [34] (version: 1.0.2) is used, for its implementation of *Bene*. The *ARTIVA* source code is publicly available as an `R` package with the same name ([35], version: 1.2.3). The implementations of algorithms {*TVDBN-0, TVDBN-bino-hard, TVDBN-bino-soft, TVDBN-exp-hard, TVDBN-exp-soft*}^2^ are available in `R` package EDISON ([36], version: 1.1.1). Experiments are performed on an Intel^®^ computing server. Its configuration is provided in Section 2.2 of the supplementary document.

### 4.4 Evaluation Metrics for Comparative Study of the Learning Power

Since the true networks are single network GRNs and the outputs of *TBN* and *TGS* are time-varying GRNs, the output set of networks 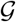 for each algorithm are converted (‘rolled up’) into an equivalent single network *G* by the following algorithm: Add a directed edge from *v_i_* to *v_j_* in *G* if there exists at least one edge from *v_i_*_*t_p_* to *v_j_*_*t*_(*p*+1)_ in 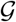 for any 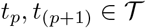. For DREAM3 synthetic datasets, self-loops (if any) are removed from the rolled network *G* since the true networks do not contain self-loops. On the other hand, for Dm datasets, self-loops (if any) are not removed from *G* as it is not known a priori whether the true network contains any self-loop or not. The metrics used to evaluate correctness of each predicted (rolled) network w.r.t. the corresponding true network are described in Section 4.7 of the supplementary document.

### 4.5 Learning From Dataset Ds10n

For Ds10n, *TGS+* outperforms every other algorithm w.r.t. every metric (Table 3a). *TBN* also achieves the highest TP but at the cost of the worst FP. Amongst *TBN* and *TGS*, it is found that *TGS* is faster, which is expected, since the regulator search space for each gene, in case of *TGS*, is monotonically smaller than that of *TBN*. But the interesting observation is that *TGS*, being a heuristic based approximate search algorithm, performs competitively with *TBN*, an exhaustive search algorithm, in every metric of learning power as well. The reason behind that is explained by the fact that the *CLR* step, in *TGS*, captures 7 out of 10 true edges even from this noisy dataset. The high TPR of *CLR* step is utilized by the downstream *Bene* step. *Bene* identifies at least as many true edges as that of *TBN*; at the same time, *Bene* avoids to search for as many potential false edges as possible. This reasoning is supported by another fact that *TGS* suffers from much less FP than *TBN*. Still, *TGS*’s FP is higher than those of the existing algorithms. This issue is taken care of, by the *ARACNE* step, in *TGS+*. Moreover, by producing a sparser mutual information matrix, *ARACNE* helps *TGS+* to have a smaller search space and, in turn, a shorter runtime.

**TABLE 3.**
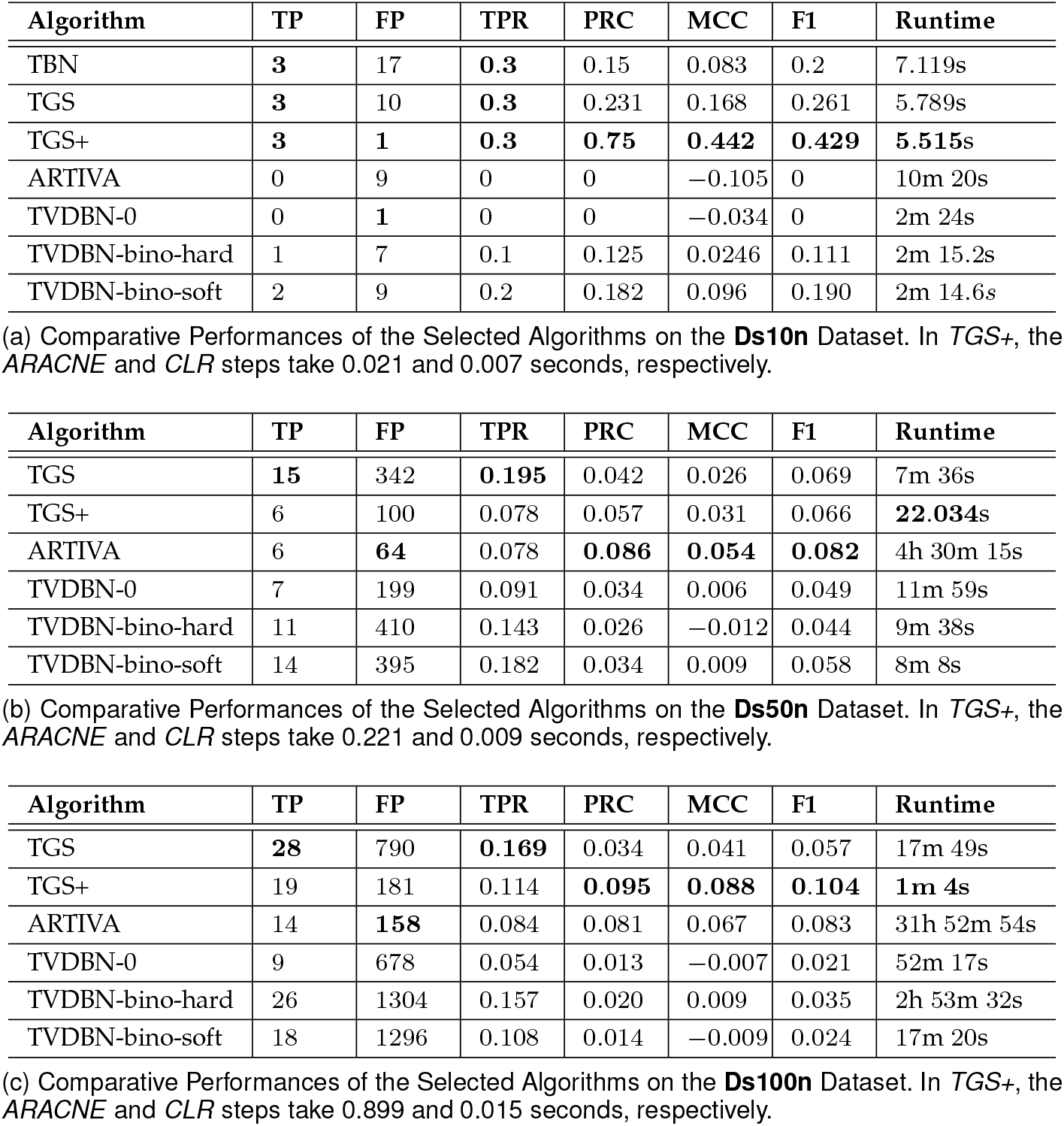
Comparative Performances of the Selected Algorithms on the DREAM3 Datasets. TP = True Positive, FP = False Positive, TPR = True Positive Rate, PRC = Precision, MCC = Matthew’s Correlation Coefficient, F1 = F1 score. The numerical values are rounded off to three decimal places. For each column, the best value(s) is boldfaced.

### 4.6 Learning From Datasets Ds50n and Ds100n

Due to *Bene*’s main memory requirement of 2^(*V*+2)^ Bytes [11], both *TBN* and *TGS* have the same inherent exponential memory requirement. In theory, that should enable them to learn a network with *V* ≤ 32 with a 31 GB main memory, since 2^(32+2)^ Bytes = 16 GB < 31 GB. But it is found empirically that the bnstruct’s implementation of *Bene* can learn a network with *V* ≤ 15 with that configuration, without any segmentation fault. Therefore, the max fan-in variant of *TGS* is employed for Ds50n and Ds100n with *M_f_* = 14, since that would restrict each atomic network learning problem to a maximum of 15 nodes (1 regulatee and a maximum of 14 candidate regulators). But *TBN* does not have any such provision and hence can not be applied on these datasets.

In this study, *TGS* and *ARTIVA* consistently produce the highest TP and the lowest FP, respectively (Table 3b, 3c). Here, *TGS+* provides a middle ground. It produces monotonically higher TPs than those of *ARTIVA*, while maintaining competitive FPs. On the other hand, the *ARACNE* step in *TGS+*, causes TPR to decline by 9% and 33% w.r.t. TGS, for Ds50n and Ds100n, respectively. But there is 0% decline for Ds10n. This observation can be explained by the fact that around 39% of true edges are feed-forward edges for Ds50n and Ds100n, whereas that is only 10% in case of Ds10n.

Another major concern with {*ARTIVA, TVDBN-0, TVDBN-bino-hard*} is the runtime. For example, *ARTIVA* takes around 32 hours to reconstruct 100-gene GRNs, which is certainly a bottleneck for its application in reconstructing human genome-scale GRNs. In comparison, *TGS* and *TGS+* consume only 18 minutes and 1 minute, respectively. Only *TVDBN-bino-soft* is able to provide a competitive runtime. However, it is consistently outperformed in {PRC, MCC, F1} by both *TGS* and *TGS+*. Moreover, the runtime of {TGS, *TGS*+} grow almost linearly as the number of genes grow (Figure 5). These observations indicate that *TGS* and *TGS+* are substantially more suitable for reconstructing large-scale GRNs than the alternative algorithms.

**Fig. 5.**
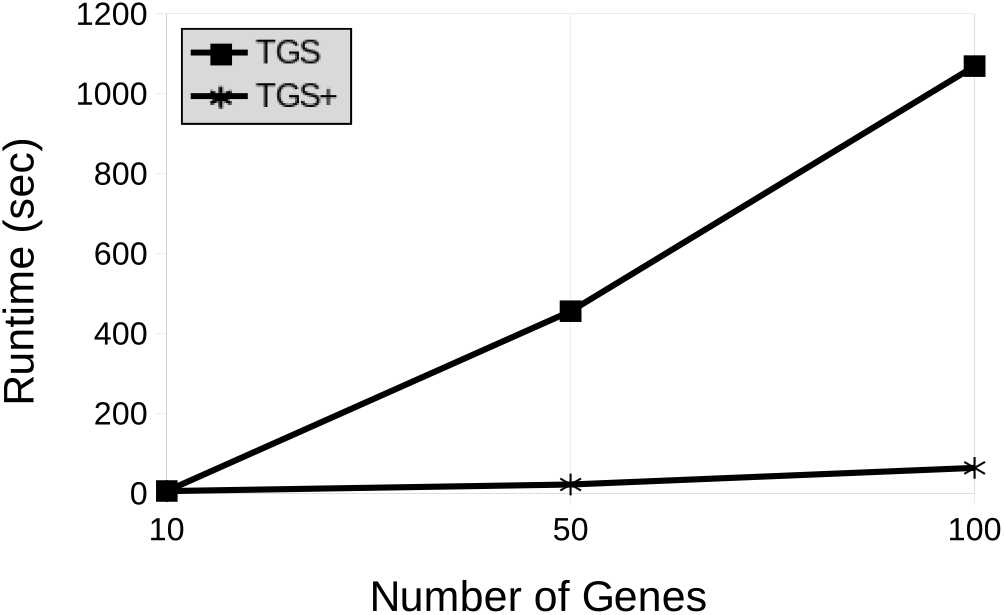
The Runtime of the *TGS* and *TGS+* Algorithms w.r.t. the Number of Genes in the Synthetic Datasets (Table 2).

### 4.7 Effect of Noise on Learning Power and Speed

*TGS* is evaluated on all noisy and noiseless datasets with different number of genes. From Figures 6 and 7, it can be observed that the presence of noise negatively impacts runtime and precision. This observation can be explained by analysing the effect of noise on the *CLR* step (Table 4). In the absence of noise, the *CLR* step can eliminate more number of potential false regulators from the candidate set of regulators of each regulatee, resulting in smaller and more precise shortlist of candidate regulators. That in turn, improves precision and speed of the overall algorithm. The effect of noise on the performance of *TGS+* is discussed in Section 4.8 of the supplementary document.

**Fig. 6.**
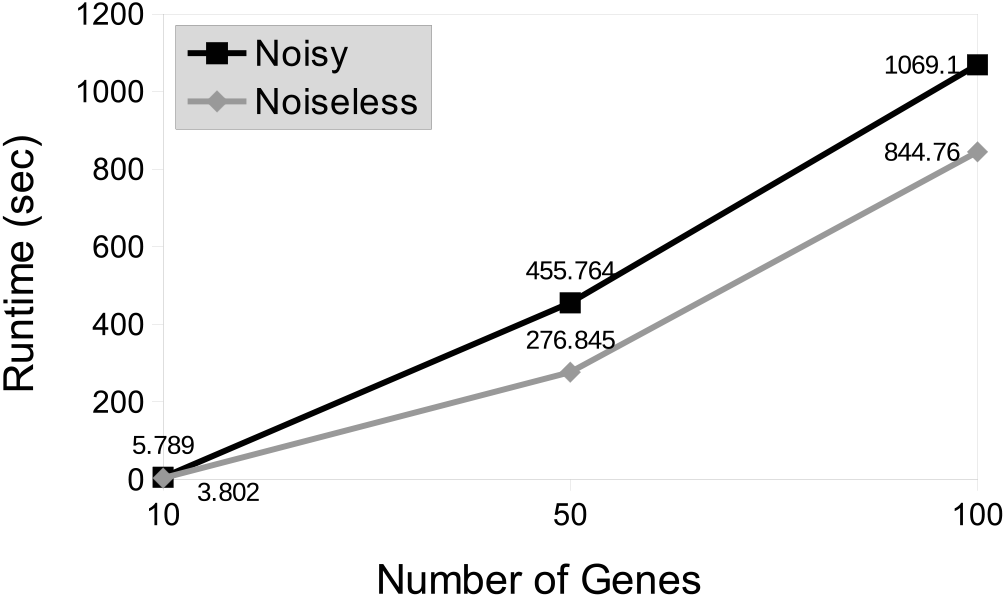
The Runtime of the TGS Algorithm w.r.t. the Number of Genes in the Input Synthetic Datasets (Table 2). The black and grey lines represent noisy and noiseless versions of the datasets, respectively.

**Fig. 7.**
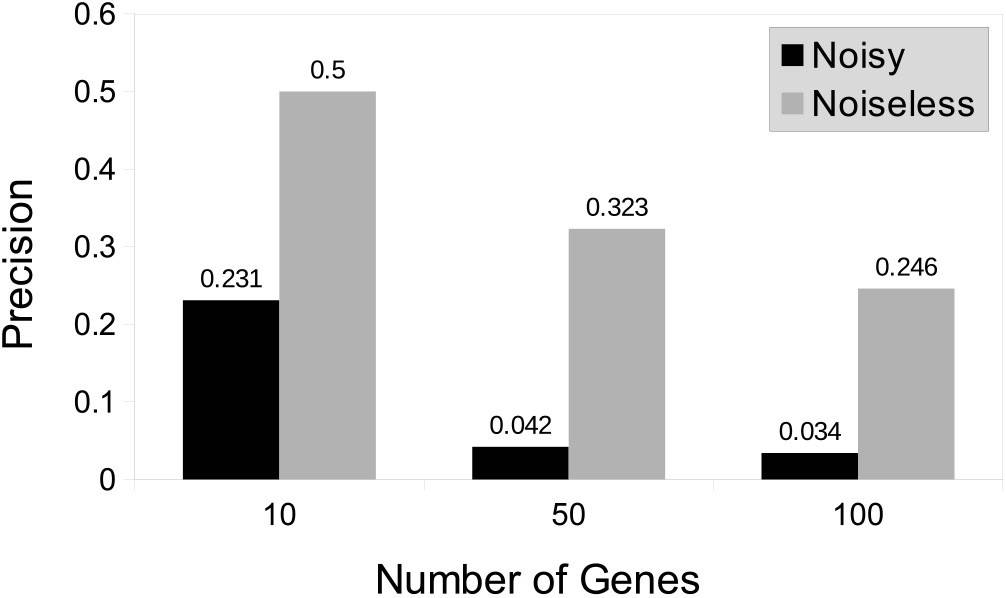
Precision of the *TGS* Algorithm w.r.t. the Number of Genes in the Datasets. The black and grey bars represent noisy and noiseless versions of the datasets, respectively. The *2L.wt* algorithm is used for data discretization.

**TABLE 4.** Maximum Number of Neighbours a Gene has in the *CLR* Network. Algorithm *2L.wt* is used for data discretization.

| Total Number of Genes | Noiseless Dataset | Noisy Dataset |
| --- | --- | --- |
| 10 | 4 | 7 |
| 50 | 24 | 33 |
| 100 | 43 | 84 |

### 4.8 Learning from the DmLc3 Datasets

For each sub-dataset of the DmLc3 dataset, all time points belong to a single time series. However, *TGS* requires multiple time series as input. Hence, the time points are divided into multiple groups and each group is considered as a distinct time series. For example, DmLcE is originally comprised of 30 time points belonging to a single time series. These time points are used to generate 5 time series having 6 time points each, with the following strategy: time points 1 − 5 are assigned to time series 1 − 5, respectively; similarly, time points 6 − 10 are assigned to time series 1 − 5, respectively; and so on. In short, the *i^th^* time point is assigned to the (*i* mod *S*)^*th*^ time series, where *i* varies from 1 to the total number of time points in the original time series and *S* equals the number of time series to be generated. This strategy ensures that the replicates at each newly created time point are consecutive time points in the original time series. For example, there are 5 replicates at the newly created first time point. It comprises of the first time points of every newly created time series. Therefore, these replicates are time points 1 − 5 in the original time series. The same single to multiple time series conversion strategy is followed for datasets DmLcL, DmLcP and DmLcA (Table 5).

**TABLE 5.**
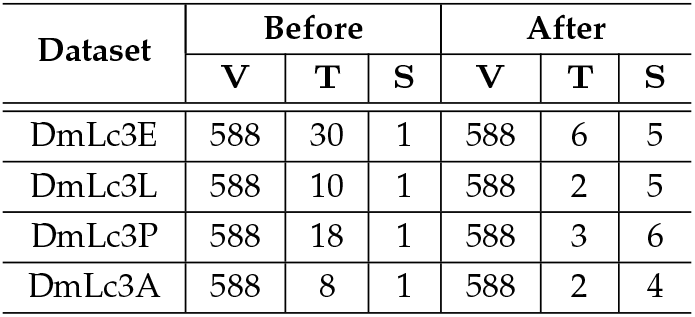
A Summary of the DmLc3 Datasets before and after the Conversion from Single to Multiple Time Series. The datasets, after conversion, are used for experimentation. Here, *V* = number of genes, *T* = number of time points and *S* = number of time series in a given dataset.

The *TGS* algorithm is applied separately on each converted dataset. It results in reconstruction of (*T* − 1) time-varying GRN(s) (one GRN for each time interval) for each dataset, where *T* = number of time points in that dataset. Therefore, five time-varying GRNs 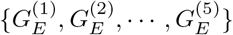 are reconstructed from DmLc3E; also a single time-invariant GRN *G_E_* is generated by rolling up the time-varying GRNs. For DmLc3L, one time-varying GRN *G_L_* is reconstructed; since there is only one time-varying GRN, there is no need to roll it up. For DmLc3P, two time-varying GRNs 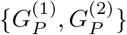 are reconstructed; also a single time-invariant GRN *G_P_* is generated by rolling up the time-varying GRNs. For DmLc3A, one time-varying GRN *G_A_* is reconstructed; again since there is only one time-varying GRN, there is no need to roll it up. The accuracy of all the reconstructed time-varying GRNs and the time taken to reconstruct them are studied in the subsequent sections.

#### 4.8.1 Study of Learning Power with the DmLc3 Datasets

The study is conducted in two steps. In the first step, a coarse-grained analysis is performed at the network level. It analyses structural properties of the predicted GRNs. In the second step, a fine-grained analysis is performed at the edge level. It attempts to evaluate the biological relevance of the predicted edges. Since, the true GRNs are not known, the evaluation is performed against the existing biological knowledge. The outcome of the coarse-grained analysis is discussed in Section 4.9 of the supplementary document. The outcome of the fine-grained analysis is described below.

**Fine-grained Analysis:** For the fine-grained analysis, a subset of 25 genes are chosen, which are known to produce TFs in Dm. This subset is generated by intersecting the set of genes in the DmLc3 dataset with the set of known TF-coding Dm genes used in Marbach et al. [37]. Then for each gene in the subset, two questions are posed:

Q1. Whether the given gene is predicted to play any regulatory role in the development stage(s) where it is known to do so? This question is answered by checking whether the gene has at least one regulatee in the predicted GRN(s) specific to that stage(s).

Q2. If answer to Q1 is yes, then does the given gene regulate any of its known regulatees (if any) in the predicted GRN(s)? This question is inapplicable when answer to Q1 is no.

For this analysis, known regulatory stages and known regulatees, if any, of each concerned gene are retrieved from TRANSFAC Public Database version 7.0 [38], which is claimed to be the gold standard in the area of transcriptional regulation [39]. This analysis finds a number of biological supports for the predicted GRNs. Some of the findings are discussed below.

**prd:** Gene ‘prd’ is known to have a positive cell specificity in the E stage. It is also known that ‘prd’ participates in the regulation of anterior-posterior segmentation of the embryo. ***TGS::*** In agreement with the prior knowledge, ‘prd’ is predicted to have the maximum number (i.e. 8) of regulatees in *G_E_*, whereas it has 3 regulatees in *G_P_*, and does not have any regulatee in *G_L_* and *G_A_*. Additionally, in *G_E_*, ‘prd’ regulates ‘eve’, which is a known regulatee of ‘prd’. ***TGS+::*** Similar to *TGS*, ‘prd’ is predicted to have regulatees only in *G_E_* and *G_P_*. However, *TGS+* misses the true edge from ‘prd’ to ‘eve’, unlike *TGS*.

**bcd:** Similar to ‘prd’, ‘bcd’ is known to be a major regulator in the anterior-posterior axis formation of the embryo. ***TGS::*** The prediction is consistent with the knowledge as ‘bcd’ has 9 regulatees in *G_E_* and does not possess any regulatee in either of {*G_L_, G_P_, G_A_*}. ‘bcd’ is also known to be a concentration-dependent regulator of ‘eve’. But there is no directed edge from ‘bcd’ to ‘eve’ in *G_E_*. It might be a true negative prediction, if the regulation did not happen during the data collection period, owing to the absence of the required concentration level. ***TGS+::*** ‘bcd’ has 3 and 2 regulatees in *G_E_* and *G_P_*, respectively. Like *TGS*, it does not possess any regulatees in other stages.

Some more genes, like - ‘tll’, ‘dl’, ‘ftz.f1’ and ‘Trl’, are reported, in literature, to play regulatory roles in the E stage. ***TGS::*** In accordance with the literature, they are found to have no regulatee in any predicted networks except in *G_E_*. Moreover, ‘Trl’ produces a very abundant nuclear protein, known as the GAGA protein. This protein has implications in the transcriptions of numerous Dm genes by either directly binding to the regulatee gene’s TF binding site or by allowing the regulatee gene to open up for transcription via modification of the chromatin configuration around it [40]. This implication is also found in *G_E_*, where ‘Trl’ is predicted to have directed paths to a total of 538 genes (downstream regulatees), in spite of having directed edges to only 2 genes (direct regulatees). ***TGS+::*** Like *TGS*, the aforementioned genes have regulatees only in *G_E_*. ‘ftz.f1’ is the only exception; it does not possess any regulatees in any of the predicted GRNs.

Another interesting prediction is found for gene ‘Antp’. Appel et al. [41] propose that, in some type of neuronal cells, the regulator proteins of ‘Antp’ compete to regulate it with the protein encoded by ‘Antp’ itself; in other words, ‘Antp’ appears to auto-regulate itself. But, in none of the GRNs, predicted by *TGS*, ‘Antp’ has a self-loop. However in *G_E_*, it does have two feedback loops (directed paths to itself), each of length 2, through genes ‘odd’ and ‘CG12896’, respectively. Whether the auto-regulation of ‘Antp’ happens through a self-loop or multi-hop feedback loops opens up an intriguing question in experimental biology.

For the complete set of findings of the fine-grained analysis, please see Table 3.1 of the supplementary document.

#### 4.8.2 Study of Learning Speed with the DmLc3 Datasets

The comparative study of DmLc3 datasets reveals that the runtime of the *TGS* and *TGS+* algorithms increase with the value of *T*, when *V* is fixed (Figure 8). This finding is consistent with the time complexity expressions of *TGS* (Equation 1) and *TGS+* (Section 4.4, supplementary document).

**Fig. 8.**
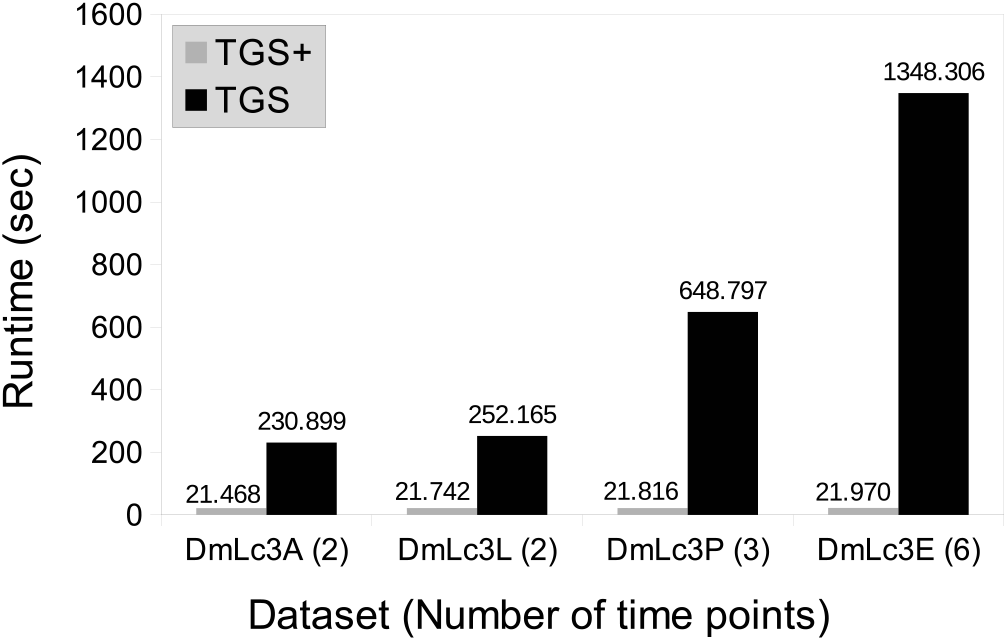
The Runtime of the *TGS* and *TGS+* algorithms w.r.t. DmLc3 datasets. These datasets (Table 5) have the same number of genes (*V* = 588). But they vary in the number of time points (*T*). It is evident that the runtime strictly increases with *T* when *V* is fixed, which is consistent with the time complexity expressions of *TGS* (Equation 1) and *TGS+* (Section 4.4, supplementary document).

An additional experiment is performed to examine whether *TGS* and *TGS+* can scale up to larger datasets, than the DmLc3 datasets, in a reasonable time frame. For this experiment, the expression levels of all 4028 genes of the DmLc dataset during the *E* stage are used. The *E* stage consists of 30 time points belonging to a single time series. They are converted to 15 time series, each consisting of 2 time points using the strategy discussed in Section 4.8. The resultant dataset is named DmLcE. *TGS* is able to scale up to DmLcE, reconstructing a GRN with 3318 directed edges, in 47 minutes. *TGS+* performs the same task, slightly faster (in 44 minutes), reconstructing a marginally sparser GRN (with 3312 directed edges).

Detailed guidelines of how to reproduce the results, presented in this section, are provided in the supplementary document.

## 5 Summary and Future Work

In this paper, a novel algorithm, namely *TGS*, is proposed to reconstruct time-varying GRNs from a time series gene expression dataset. *TGS* assumes that there are multiple time series and no missing values in the dataset.

*TGS* employs a two-step learning strategy. In the first step, for each target gene, a shortlist of its potential regulators is inferred. In the final step, these shortlisted candidates are thoroughly evaluated to identify the true regulators among them. Moreover, the temporal sequence of the regulatory events is learnt from the data.

The novelty of the *TGS* algorithm is two-fold: (A) flexibility and (B) time-efficiency. Its flexible framework allows time-varying GRNs to be learnt independently of each other. *TGS* learns every GRN structure in a data-driven manner, without imposing any structural constraints.

However, an existing algorithm, namely *ARTIVA*, provides a similarly flexible framework [4]. The only challenge with *ARTIVA* is its substantial runtime. That makes *ARTIVA’s* application prohibitive with high-throughput datasets. *TGS*, on the other hand, is able to offer the same flexibility in a significantly more time-efficient manner.

It needs to be noted that the flexibility comes at a cost. Learning each time interval specific GRN, independently, demands sufficient number of measurements per time point, for each gene. Empirically, it is seen that *TGS* needs at least 4 replicates per time point, for each gene, to reconstruct meaningful GRNs. If the number of measurements is low, then *TGS*, like ARTIVA, may lead to overfitting [5]. For such datasets, it is preferable to use less flexible algorithms, e.g., Dondelinger et al. [5], which learns the GRNs jointly, by sharing information across time intervals.

However, the joint learning algorithms perform well, only when changes in true GRNs, across time intervals, are gradual (‘smoothly time-varying’ GRNs). That is not the case, always. For example, introduction of a drug, during treatment, can result in a drastic change [5]. On the other hand, *TGS*’s framework is compatible with any dataset, regardless of how smooth the underlying changes are.

Moreover, *TGS* provides such flexibility and time-efficiency, without losing its accuracy. It consistently outperforms *ARTIVA* in true positive detection (sensitivity), given three benchmark, realistically simulated, datasets. On the other hand, *ARTIVA* performs better, in false positive rejection. Hence, a less sensitive but more precise variant of *TGS*, namely *TGS*+, is proposed. *TGS+* is comparable to *ARTIVA*, in false positive rejection, while still being more sensitive.

Nevertheless, there are scopes for improvement. One limitation of *TGS* is the need to discretize data. The reason behind that is *TGS* uses BIC score to determine the best GRN structure and BIC scoring function requires the data to be discretized. Two of the ways the issue can be resolved are: (a) by using a scoring function that does not require discretized data, like in Grzegorczyk et al. [3], and (b) by developing a regression based structure learning strategy, e.g., Lèbre et al. [4], because regression problems are inherently compatible with continuous data.

Another area for improvement is the temporal resolution of the shortlisting strategy. In *TGS*, the shortlist of candidate regulators, for each gene, is time-invarint. Therefore, the regulators, which are active over a small number of time intervals, may not get shortlisted. To amend this issue, time interval specific shortlists can be generated, for each gene.

One more area for improvement is the utilization of existing domain knowledge, apart from gene expression data. The prediction accuracy of *TGS* can be further improved by integrating known information, if any, about the concerned system, like - protein expression profiles, protein-DNA interaction data, protein-protein interaction data, DNA-binding sequences etc., e.g., Jain et al. [42].

However, the experiments with the large datasets help the authors to identify the Achilles’ heel of *TGS*. It is the fact that its main memory requirement grows exponentially with the number of genes (and in turn number of candidate regulators for each gene) in the datasets. In the current implementation of *TGS*, maximum number of candidate regulators is restricted to 14 for each gene, to avoid this issue. But relaxing this restriction is an important challenge since the actual number of regulators for a gene is not known a priori.

The reason behind such astronomical memory requirement is that *Bene* and related Bayesian Network structure learning algorithms need to compute and store the global conditional probability table [11] in main memory. Some researchers are exploring efficient ways to distribute this task and storage across multiple computing nodes using distributed computing strategies, e.g., Jahnsson et al. [43]. Mending this gap can be considered a worthwhile challenge. Convergence of affordable high-throughput gene expression measurement technologies with accurate and scalable GRN reconstruction methodologies will be a valuable achievement. It will help in improving our understanding of disease progression and life in general through the lens of gene regulation.

## Acknowledgements

SP and ARK received MHRD Fellowships from IITG during this work.

---

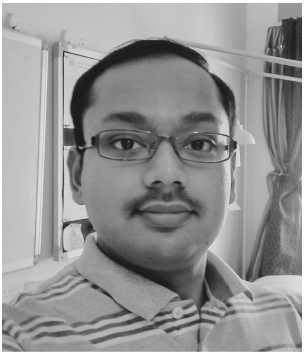

**Saptarshi Pyne** is a PhD student in Dr. Ashish Anand’s research group. His research area is temporal progression modelling of biological systems. Saptarshi believes in a future where biomolecular signals are measured in-vivo and analysed in near real time. **Website:** http://iitg.ac.in/stud/p.saptarshi/

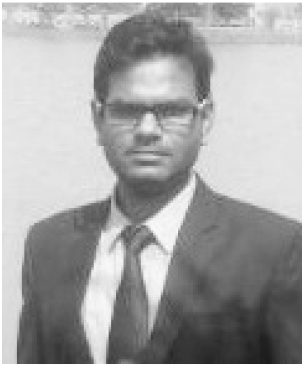

**Alok Ranjan Kumar** wrote his MTech thesis, titled ‘Structure learning of gene regulatory network from large-scale (high-dimensional) time-series gene expression data’, under the supervision of Dr. Ashish Anand. Alok is a graduate trainee engineer at Siemens Industry Software (India) Pvt. Ltd. **LinkedIn:** https://www.linkedin.com/in/alok-kumar-4039/.

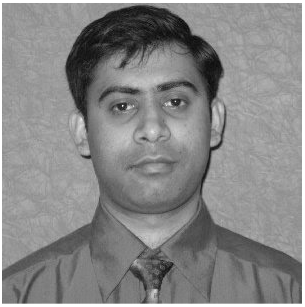

**Ashish Anand** is an assistant professor at the Department of Computer Science and Engineering, Indian Institute of Technology Guwahati (IITG), India. Prior to joining IITG, he was a member of the European Consortium, BaSyS-Bio at the Systems Biology Lab, Institut Pasteur, Paris. His main research area is temporal progression modelling of biological systems. **Website:** http://iitg.ac.in/anand.ashish/

## Footnotes

1. An MI network is an undirected graph where two nodes are connected if and only if their pairwise MI is statistically significant. The significance threshold is either user defined or programmatically computed by the network inference algorithm itself.

2. The implementations of *TVDBN-exp-hard* and *TVDBN-exp-soft* crashed in all the experiments. In a personal communication with the authors, the author-cum-maintainer of the package hinted at a potential bug, that might be fixed in a future version of the package. Therefore, the results of *TVDBN-exp-hard* and *TVDBN-exp-soft* are not reported in this paper.

